# A minimal activator-inhibitor-repressor model of the hepatic circadian clock

**DOI:** 10.1101/2025.08.08.669290

**Authors:** Pauline Delpierre, Marc Lefranc

## Abstract

Circadian clocks rely on gene regulation networks which generate periodic biochemical oscillations informing our cells about the time of the day. Mathematical modeling has been effective to describe the dynamics of the multiple intertwined feedback loops making up circadian clocks, however it is often delicate to adapt the complexity of the model to the question addressed and to the data available. Traditionally, two main modeling approaches have been followed, using either comprehensive models recapitulating most molecular actors involved, or minimal qualitative models highlighting the core mechanisms. However, analyzing the behavior of large models may be difficult, and small models often lack predictive power, questioning their relevance. Through a systematic reduction of a more complex model, we obtain a simple three-gene clock model, featuring the activator *Bmal1*, the repressor *Reverb* and the inhibitor *Cry*, that accurately describes the corresponding temporal expression profiles for the mouse hepatic clock. We characterize this model by carrying out a sensitivity analysis for its limit cycle, as well as by computing phase response curves for the different possible inputs. Predictions from the model are compatible with a number of synchronizing mechanisms from the literature.

**SIGNIFICANCE:** Living systems adapt to the day/night cycle thanks to cellular clocks, which track the time of the day and orchestrate physiological processes throughout the 24 hours. The dynamics of these clocks is complex, due to the interaction of intertwined feedback loops generating the biochemical oscillations and ensuring their synchronization. Mathematical modeling has proved useful to unravel this complexity, however there is usually a difficult choice to be made between comprehensive and minimal models, with opposite strengths and weaknesses. Here, we propose a simple activator-inhibitor-repressor model reproducing surprinsingly well experimental data from mouse livers, whose analysis casts light on the roles of the main actors of the mammalian circadian clock.

## INTRODUCTION

Most living organisms on Earth are subjected to an alternation of days and nights due to Earth rotation. To anticipate the associated daily changes in their environment, they have evolved a circadian clock, a genetic oscillator where genes and proteins regulate each other so as to generate oscillations with a period of approximately 24 hours (1, 2). The regulation networks that underlie circadian clocks feature many interlocked feedback loops whose collective dynamics is complex and defies mere intuition. Hence, their understanding calls for quantitative approaches that can integrate the concurrent interactions generating the clock oscillations. In particular, mathematical modeling has been successful in disentangling the complexity of the circadian clock (3–9).

A typical mathematical model of the circadian clock recapitulates molecular actors and interactions that have been shown experimentally to contribute to clock function, or a subset thereof. It describes the rate at which the activities of the involved mRNA and protein molecules vary in time, based on our knowledge of the underlying biochemical reactions. Not all kinetic constants are known a priori, but their values can be adjusted by fitting the model to experimental data. For a given system, different models may be elaborated depending on the context. Indeed, a mathematical model depends on (1) the system considered, (2) the question addressed, and (3) the data available. Different clock actors have generally different functions, and may play a major or a minor role depending on the specific behavior that is investigated.

A mathematical model can serve several purposes. First, it can be used to check the consistency of the various interactions assumed. Reproducing complex oscillations such as those generated by a circadian clock is challenging, as changing a single regulation can often ruin the agreement or even suppress oscillations. Therefore, a good agreement between a circadian clock model and the data is generally satisfying (see, e.g. (3)), even if it is difficult to tell apart molecular mechanisms that similar dynamical effects. Models which are contradicted by experimental facts are still useful, as this guides us toward searching for other mechanisms not yet taken into account.

Mathematical models also cast light on the interplay of the actors considered, and their relative roles in a mechanism. Kinetic constants are easily modified in numerical experiments where various interactions are switched on or off, testing the importance of actors or regulations. In particular, techniques such as sensitivity analysis indicate the kinetic constants on which a specific behavior depends principally, allowing us to isolate the backbone of the regulation network which is relevant to the behavior studied (10–13). Assessing parameter identifiability indicates which parameters are completely constrained by the available data, and which ones have an uncertain value (14–17). Mathematical models can also be used to determine which experiments would bring the most new information about the system (18).

In the end, no mathematical model should be completely trusted until it has been extensively validated experimentally. As such, it only reflects the current knowledge of a system, but still may prove useful to guide future experimental and theoretical investigations.

Two opposite strategies may be followed to build a mathematical model. On the one hand, one may seek to include all documented actors and interactions, thus summarizing the data from the litterature. When first addressing a biological question, this may be important to ensure that no important actor is overlooked. Such models may also be necessary when quantitative predictions are required, for example when designing therapeutical protocols where delivery doses and timings must be determined precisely. However, the model may be too complex to allow identifying the key mechanisms. Moreover, the training of large models require extensive data sets, acquired in various conditions. Designing such a model when only limited data are available often leads to an under-determined model where multiple parameter sets can fit the experimental data. This is not necessarily a problem, as many system biology models have nonidentifiable parameters (19). However, the predictions of such a model are often poor outside of the original experimental context, because they depend on the specific parameter set used.

On the other hand, one may seek to design minimal models highlighting the core ingredients of a mechanism (see, e.g. (20). Such models may be obtained by gradually incoporating actors and regulations, until the desired agreement with experimental data is reached. Alternatively, as we do here, one may start from a complex model and reduce it while preserving the agreement with data (see, e.g., (21–23). The advantage of this approach is that every model ingredient is by construction necessary, even if two molecular mechanism inducing the same dynamical effects cannot be distinguished. Minimal models nicely illustrate the roles played by the key actors, but often have limited predictive power since the emphasis is not on matching the data accurately. This may affect the relevance of the mechanistic insights they provide.

In this work, we design a minimal three-gene circadian clock model that nevertheless reproduces accurately temporal expression profiles of the murine hepatic clock, both for mRNA and protein abundances. We obtain it by gradually reducing a larger model in a controlled way. Usually, molecular actors are removed from a model when a simulated knock-out has little effect on clock behavior, showing that its presence is not needed. However, another important and perhaps more relevant question about the role of a circadian actor is whether its temporal variations are essential to the clock dynamics. What happens if we find a way to gradually dampen its dynamics so as to make its abundance eventually constant, or at least its influence on other actors? In other words, are the variations of this actor a mere consequence of the clock dynamics or are they essential to generate it?

Here, we apply this strategy to a five-gene model derived from the core clock section of the Woller model (8), and eventually obtain a three-gene model where the activator *Bmal1*, the inhibitor *Cry* and the repressor *RevErb* interact to generate oscillations that reproduce very well the experimental data for these three genes, as well as a two-gene model that accounts well for the expression profiles of *Bmal1* and *RevErb* mRNA and protein products. The method of “clamping” some molecular actors has been previously used by Pett et al. (24), although they applied it to a slightly different clock model and did not use a gradual process as here. They identified a core loop comprising the *Reverb, Per* and *Cry* genes arranged in a “repressilator” motif. Del Olmo et al. (22) followed also a similar approach to describe circadian redox oscillations.

Finally, we carry out a sensitivity analysis of the model and compute phase response curves for a variety of clock inputs. This allows us to identify those inputs whose variation has the most influence on the limit cycle oscillations and on the clock phase. We find that a few inputs are suitable for synchronizing the liver clock to an alternation of its metabolic state between day and night, and that they are consistent with experimental observations.

## METHODS

Numerical simulations and parameter optimization were performed using the COPASI pathway simulator (25), as well as Python scripts utilizing its programming interface basico (26). The coefficients of variation in Tables 1 and 2, quantifying the uncertainty in the determination of a parameter, were those reported by the COPASI Parameter Estimation task.

**Table 1.** Value of the control parameters of the *Bmal1*-*Reverb-* Cry model for which the profiles of Fig. 2 have been obtained. Parameters are ranked by order of increasing uncertainty, as determined by their coefficient of variation in parameter sets fitting the data, from more identifiable to less identifiable.

| Constant | Value | CV (%) |
| --- | --- | --- |
| $t_{rb}$ | 1.24157 | 3.6 |
| $k_r$ | 0.400458 | 3.7 |
| $d_{pr}$ | 0.360599 | 4.0 |
| $t_{cb}$ | 0.823388 | 4.5 |
| $d_{mr}$ | 0.520944 | 5.4 |
| $t_{br}$ | 0.485667 | 5.5 |
| $h_{br}$ | 2.3885 | 6.0 |
| $k_c$ | 0.151206 | 6.1 |
| $d_{pc}$ | 0.150489 | 6.3 |
| $d_{mb}$ | 0.743706 | 6.4 |
| $b_b$ | 1.72303 | 6.8 |
| $h_{cr}$ | 1.24357 | 8.9 |
| $h_{rc}$ | 6.95435 | 10.1 |
| $k_b$ | 0.0424037 | 11.4 |
| $d_{pb}$ | 0.0425099 | 11.5 |
| $t_{cc}$ | 0.983016 | 11.8 |
| $h_{cc}$ | 4.7554 | 12.3 |
| $h_{rb}$ | 31.566 | 12.6 |
| $t_{rc}$ | 0.847034 | 15.4 |
| $b_r$ | 0.0341104 | 27.5 |
| $h_{cb}$ | 16.81 | 32.4 |
| $t_{cr}$ | 0.105741 | 60.1 |
| $d_{mc}$ | 2.7455 | 87.1 |
| $f_r$ | 944.095 | 88.1 |
| $f_c$ | 2000 | 1656 |
| $b_c$ | 0.0344934 | 1667 |

**Table 2.** Value of the control parameters of the two-gene *Bmal1*-*Reverb* model for which the profiles of Fig. 3 have been obtained. Parameters are ranked by order of increasing uncertainty, as determined by their coefficient of variation in parameter sets fitting the data, from more identifiable to less identifiable.

| Parameter | Value | CV (%) |
| --- | --- | --- |
| $t_{rb}$ | 1.12378 | 3.3 |
| $d_{mr}$ | 0.459495 | 6.1 |
| $k_r$ | 0.386722 | 6.6 |
| $d_{pr}$ | 0.342897 | 7.0 |
| $t_{br}$ | 0.615491 | 7.9 |
| $h_{br}$ | 2.98324 | 10.0 |
| $d_{mb}$ | 0.725878 | 11.2 |
| $b_b$ | 1.56256 | 11.7 |
| $d_{pb}$ | 0.028417 | 12.5 |
| $k_b$ | 0.0285584 | 12.6 |
| $h_{rb}$ | 50 | 24.3 |
| $b_r$ | 0.0429166 | 28.3 |
| $f_r$ | 81.0185 | 43.9 |

The various models were fitted to data sets informing them about the mRNA and protein abundances of the 5 genes involved. The mRNA dataset was the same as used in Woller et al. (8), consisting of Fourier series approximating the temporal profiles obtained by Hughes et al. from mouse livers (27). This dataset has a temporal resolution of one hour, yielding well-defined profiles that strongly constrain mathematical models, and captures well the strong non-sinusoidal character of some profiles. The protein data were whole-cell data obtained by quantitative mass spectrometry from mouse liver by Narumi et al. (28). Both datasets were normalized so that the average level of each profile is 1, making the interpretation of regulation thresholds easier. Following (8), the *Bmal1, Reverb*,*Cry, Per*, and *Ror* numerical profiles were respectively fitted to the experimental profiles of *Bmal1, Reverbα, Cry1, Per2*, and *Rorγ*, a choice which is relevant for liver physiology.

To suppress as much as possible oscillations of the expression profile of a protein, we imposed an upper bound on the protein degradation rate and carried out a parameter scan of this bound down to a very small value (e.g., 10^−5^ *h*^−1^), with a parameter estimation task achieved at each step to adapt the remaining parameters to the changes in degradation rate. Given the complexity of the parameter optimization landscape, this gradual process was required to avoid losing track of the optimum. When the agreement of the model with the data can be thus preserved, this indicates that the variations of this protein do not play an essential role in the dynamics, and that the protein can be turned into a constant parameter. Since only proteins influence the transcription of other genes, the corresponding mRNA concentration can then be removed.

To remove protein complexes as dynamical variables, we similarly imposed an increasingly smaller upper bound on the complex degradation rate, then an increasingly higher lower bound on the kinetic constants of the association/dissociation reactions, eventually ensuring that the complex concentration is always at equilibrium with the individual protein concentrations. The former can then be rewritten in terms of the latter, and the complex is no longer a dynamical variable.

The sensitivity analysis of the limit cycle was done by simulating the effect of a very small perturbation. Phase response curves for the different control parameters were computed by simulating a 10% increase in the value of the parameter for one hour and for 12 hours. Both were obtained using the basico programming interface to Python.

Unlike in many theoretical studies, we did not constrain the Hill coefficients which control how sensitively transcription of a given gene reacts to variations in its transcription factors. This allows the model to account for more complex mechanistic schemes than the mere binding of a transcription factor to a promoter, such as when chromatin remodeling and/or histone modifications are involved. For example, it was shown that the CLOCK:BMAL1 complex is a pioneer-like transcription factor (29), which induces many molecular events leading eventually to transcription initiation. And indeed, a theoretical study showed how high Hill coefficients naturally arise when ultrasensitive signaling cascades mediate the action of the transcription factor, leading to switch-like behavior (30).

## RESULTS

### Systematic reduction of the initial model

For a comprehensive description of the mammalian clock, please refer to (1). The Woller mathematical model augments the traditional core clock model with additional feedback loops featuring the AMPK and SIRT1 metabolic sensors (8). The core clock part of the model is basic compared to more elaborate models (see, e.g., (3, 7, 20)). It does not take compartmentalization into account, does not distinguish between the isoforms of a given gene (e.g., the *Per*1-3 isoforms are grouped into a single *Per* gene) and assumes that *Clock* evolves synchronously with *Bmal1* (or equivalently that *Clock* expression is constant). This leads to a five-gene model (*Bmal1, RevErb, Ror, Per, Cry*), which also takes into account complex formation, such as the dimerization of PER and CRY in the PER:CRY complex or of BMAL1 and CLOCK in the CLOCK:BMAL1 complex. This model features 12 differential equations and 53 kinetic parameters (8). Using the datasets described in the Methods section, we could obtain a perfect match for the mRNA data and a relatively good match, although less precise, for the protein data (Fig. 1).

**Figure 1.**
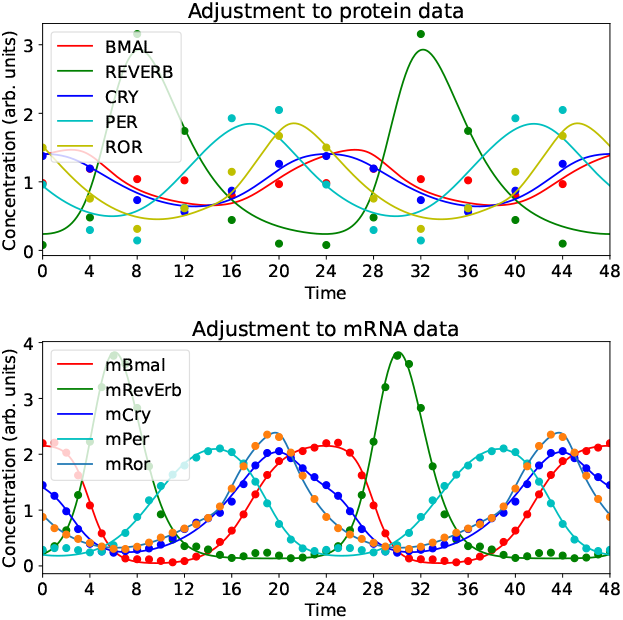
Adjusment of the five-gene model (featuring *Bmal1, Reverb, Cry, Per, Ror*) to the mRNA data of Hughes et al. (27) and the protein data of Narumi et al. (28).

Then, we proceeded to simplify this clock model, by selecting an actor and trying to gradually make its dynamics constant while simultaneously varying all other control parameters to maintain the fit to data (see Methods).

We began by eliminating the dynamics related to the PER:CRY and CLOCK:BMAL1 complexes, which could be achieved smoothly. We found that eliminating complex formation first, then complex degradation, was less robust than the converse. Indeed, the fit of the PER protein profile would then degrade, with the PER profile gradually converging to a flat, time-independent, profile.

Since reducing complex formation had suggested that the dynamics of the PER protein was not essential, we then tried to flatten it, which was easily achieved. Since the PER and CRY proteins are generally assumed to play similar roles, we investigated whether CRY expression could instead made constant. We observed that fit quality was then compromised, with the amplitude of the REVERB profile gradually decreasing. This important difference between CRY and PER may be traced back to the fact that *Cry*1, but not *Per*1-3, is repressed by REVERB*α* (31). To our knowledge, this interaction was first incorporated in a mathematical model by Relogio et al. (7) and is used in the Woller model (8). Given that CRY inhibits the induction of *Reverb* by BMAL, this suggests that the positive feedback loop comprising *Reverb* and *Cry* plays an important role in the clock dynamics.

Next, we noted that for the control parameter values used, the variations of the ROR protein had little influence on the *Bmal1* transcription rate, which essentially depended on REVERB protein abundance. Thus, we proceeded to make the ROR profile constant, which was also easily carried out.

This left us with a three-gene model based on the *Bmal1, Reverb* and *Cry* genes. As we can seen in Eqs. (1), this model comprises 6 differential equations, whose behavior is tuned by 26 control parameters. This represents an important simplification compared to the original model.

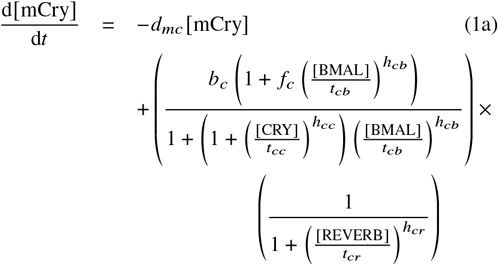

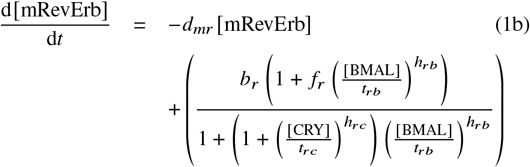

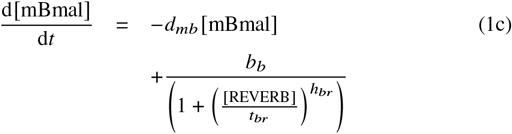

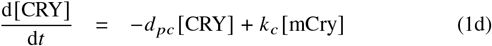

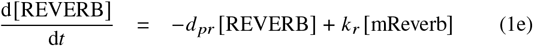

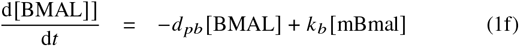

The fit of this model to the data from Hughes et al. (27) and Narumi et al. (28) led to a surprisingly good fit to the experimental data, given the simplicity of the model (Fig. 2). Remarkably, the fit to protein data was better than for the 5-gene model, presumably because optimization in this lower-dimensional parameter space is easier. The values of all control parameters are given in Table 1, ranked in order of increasing uncertainty.

**Figure 2.**
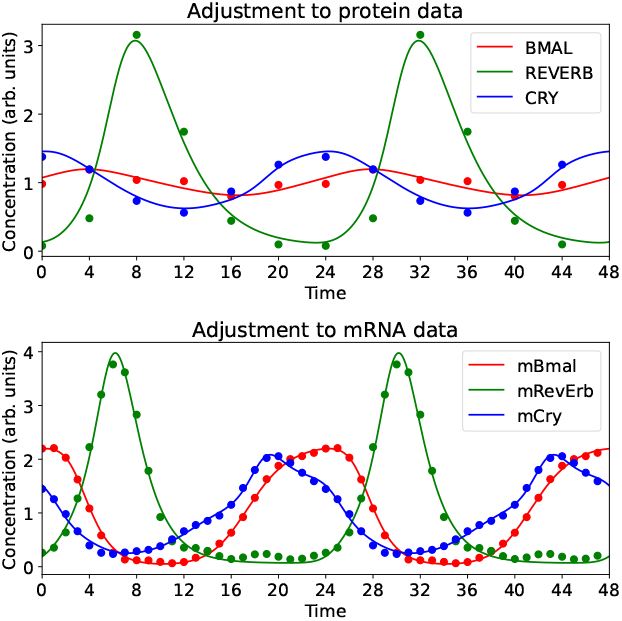
Adjusment of the three-gene model (featuring *Bmal1, Reverb*, and *Cry*) to the mRNA data of Hughes et al. (27) and the protein data of Narumi et al. (28).

One can note very high values of some Hill coefficients (see Methods) Table 1, in particular the one associated with the regulation of *Reverb* by BMAL1 (*h*_*rb*_), whose value is 31.5. This can be correlated with the contrast between a smooth BMAL1 protein profile, due to a long BMAL1 half-life, and the sharp induction of *Reverb* transcription, suggesting an almost all-or-nothing induction. Presumably, this reflects the complex cascade of regulations induced by BMAL1 (29). In contrast, the regulation of *Bmal1* by REVERB is associated with a Hill coefficient of 2.39, consistent with the fact that REVERB binds to two RRE elements in the *Bmal1* promoter (32).

Most parameters are well determined, with a coefficient of variation (CV) of 15 % at most. This ensures that the predictions of the model will be quite unambiguous. Only 5 parameters had CVs higher than 50%. We conjectured that the uncertainty of some of these parameters might be structural, especially given the astronomically high values of the CVs of *b*_*c*_ (the basal *Cry* transcription rate in the absence of transcription factors) and *f*_*c*_ (the fold change in *Cry* transcription rate upon activation). Indeed, Eqs. (1) indicate that, when *b*_*c*_ tends to zero, the transcription rate of *Cry* depends only on the product *V*_*c*_ = *b*_*c*_ *f*_*c*_. Thus, we tried to simplify the model accordingly, with *V*_*c*_ being the only parameter describing basal transcription. The fit was slightly degraded, however there remained a large uncertainty in *V*_*c*_. Thus, we kept the original model with the separate parameters for basal transcription rate and fold change. Similar observations were made for *d*_*mc*_ (*Cry* mRNA degradation rate). Together with the uncertainty in the threshold *t*_*cr*_ (threshold of *Cry* regulation by REVERB), this suggested a less important role for *Cry* than for the other two genes.

To check whether this model could be simplified further, we tried removing each of the three remaining genes in turn, beginning with *Cry* since its influence on profile fitting seemed to be weaker. This could be done easily, yielding a two-gene model based on *Bmal1* and *Reverb* (Eqs. (2)). The best-fitting parameters given by Table 2 were relatively well determined except for *b*_*r*_ and *f*_*r*_.

This model reproduces nicely the experimental data, as Fig. 3 shows, although not as well as with the three-gene model used in Fig. 2. This is an interesting result suggesting that the BMAL-REVERB negative feedback loop plays an important role in generating and shaping the clock oscillations.

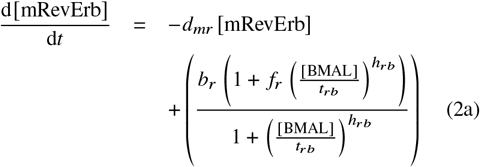

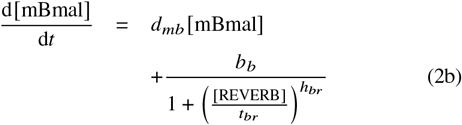

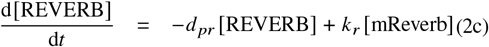

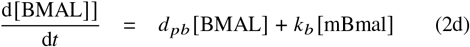

**Figure 3.**
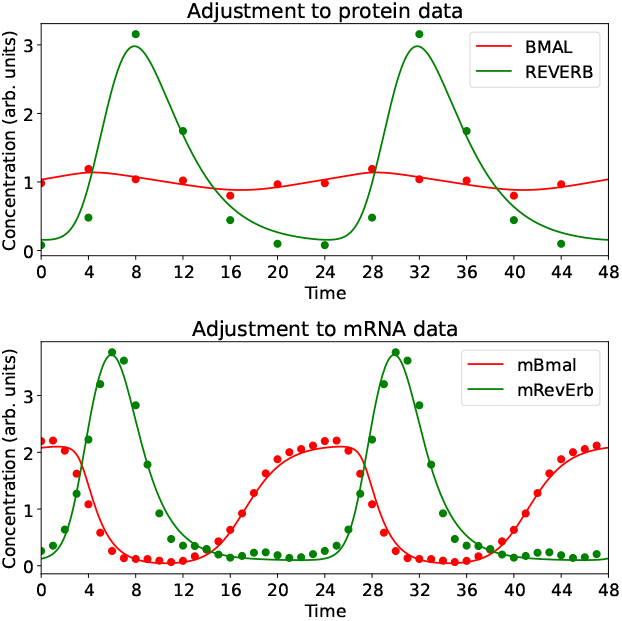
Adjusment of the two-gene model (featuring *Bmal1*, and *Reverb*) to the mRNA data of Hughes et al. (27) and the protein data of Narumi et al. (28).

Alternatively, gradually making the BMAL1 profile constant strongly degraded the fit without losing oscillations, suggesting that variations in BMAL1 activity are important to shape the circadian oscillations as we observe them. Finally, dampening the dynamics of REVERB while trying to maintain fit quality totally abolished oscillations, highlighting the key role played by the dynamics of REVERB in driving the oscillations of the circadian clock.

Given the excellent fit obtained with the three-gene model and to better identify the relative roles of the different genes, since *Bmal1, Reverb* and *Cry* have each a different mode of action, we went on to further analyse this three-gene model.

### A simple Activator-Inhibitor-Repressor (AIR) model

The *Bmal1*-*Reverb*-*Cry* model has a remarkable structure where each of the three genes plays a distinct role, influencing the other genes in its own way. BMAL1 activates *Reverb* and *Cry*; REVERB represses *Bmal1* and *Cry*; CRY inhibits the activity of BMAL1 when acting on *Reverb* and *Cry* (Fig. 4). Accordingly, we call this mathematical model the Activator-Inhibitor-Repressor (AIR) model, considering that such a separation of roles may stem from general design principles.

**Figure 4.**
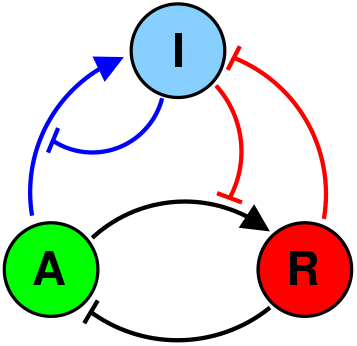
Schematic diagram of the AIR model, showing the interactions between the activator *Bmal1* (A), the repressor *Reverb* (R) and the inhibitor *Cry* (I). The three main feedback loops in this model are shown in different colors. These are the A-R (BMAL1-REVERB) negative feedback loop (black), the negative feedback loop of I (CRY) upon itself (generally described as the CLOCK:BMAL1-PER:CRY feedback loop, blue), and the less discussed but nevertheless important positive feedback loop I-R (REVERB-CRY, red).

Interestingly, the AIR model incorporates the two main negative feedback loops that have been identified in the mammalian clock (Fig. 4). The first loop links BMAL1 (representing CLOCK:BMAL1) and REVERB (drawn in black in Fig. 4), and plays an important role in driving the oscillations as demonstrated by Fig. 3. The second loop involves BMAL1 and CRY (representing PER:CRY) and is drawn in blue in Fig. 4. Stricto sensu, it describes a negative feedback of CRY on itself mediated by BMAL1, as CRY does not influence the transcription of *Bmal1*. In addition, there is a lesser known positive feedback loop formed by CRY and REVERB (drawn in red in Fig. 4), which influence each other negatively (CRY inhibits the induction of *Reverb* and REVERB represses *Cry*). As noted above, the presence of this loop is important for fitting the model to the experimental data.

In the following, we analyse the sensitivity of the three-gene model to two different types of perturbations. First, how does the limit cycle (the trajectory in state space) deviate when a parameter is changed? Second, what is the phase shift experienced when the system returns to the limit cycle solution after a transient perturbation has been applied?

### Sensivity analysis of the AIR limit cycle

We first analyzed how the limit cycle associated with oscillations in the three clock genes varies with control parameters. To this aim, we computed scaled sensitivities to variations in control parameters. More precisely, we evaluated the following quantities

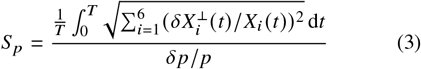

where the numerator features the average over the entire circadian cycle of the relative transverse displacement for each phase of the limit cycle in the six-dimensional state space, resulting from a variation *δp* of the control parameter *p*. The *X*_*i*_ (*t*) are the values of the 6 variables at circadian time *t*. The 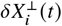 are their variations perpendicular to the limit cycle, defined by 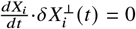, thus characterizing changes in amplitude rather than in phase. The value of *S* _*p*_ indicates by how many percent this average scaled displacement varies for a variation of the parameter of 1%. Table 3 shows the computed sensitivities larger than 1.5, from higher to lower.

**Table 3.**
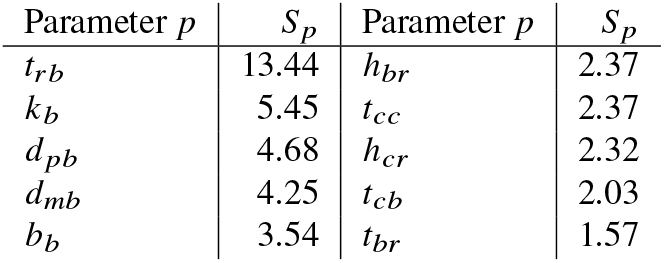
Sensitivities of the limit cycle to variations of the indicated parameters, as defined by Eq. (3). Only sensitivities larger than 1.5 have been shown.

| Parameter $p$ | $S_p$ | Parameter $p$ | $S_p$ |
| --- | --- | --- | --- |
| $t_{rb}$ | 13.44 | $h_{br}$ | 2.37 |
| $k_b$ | 5.45 | $t_{cc}$ | 2.37 |
| $d_{pb}$ | 4.68 | $h_{cr}$ | 2.32 |
| $d_{mb}$ | 4.25 | $t_{cb}$ | 2.03 |
| $b_b$ | 3.54 | $t_{br}$ | 1.57 |

The control parameter yielding the maximal variation of the limit cycle is *t*_*rb*_, the concentration of BMAL1 at which *Reverb* is half-activated. Physiologically, this parameter may for example vary when the activity of CLOCK:BMAL1 as a transcription factor is modulated by NAD+-dependent SIRT1 (33).

Then we find in Table 3 a series of kinetic constants eventually governing the abundance of the BMAL1 transcription factor: *k*_*b*_, *d*_*mb*_, *d* _*pb*_ and *b*_*b*_, respectively controlling the translation of *Bmal1*, the stabilities of *Bmal1* mRNA and protein, as well as the unrepressed expression of *Bmal1*. This suggests that *Bmal1* expression and factors that control it play a key role in driving the amplitude of clock oscillations. This is consistent with experimental observations that PGC1*α*-mediated induction of *Bmal1* has a strong effect on clock amplitude (9, 34). The remaining parameters play less important roles.

### Phase response curves of the AIR model

Another important piece of information is the phase change experienced when a transient perturbation of given duration is applied, measured after the system is allowed to return to its nominal behavior. This information is summarized by the phase response curve (PRC), which indicates the phase change depending on the time when the perturbation is applied.

The PRC is also key to understanding how a circadian clock synchronizes to an external cycle (a.k.a. a zeitgeber). Synchronization occurs when the induced phase change compensates exactly the mismatch between the periods of the clock and of the zeitgeber (35). By studing the phase response curve associated with the modulation of a given parameter, we may assess the suitability of the associated input to be a good synchronizer. Input mechanisms for which the PRC spans a wide range of phase shifts will generally be more adequate for resetting the clock and synchronize it to the external cycle. However, the shape of the PRC is also important (36).

Although our model is a free-running oscillator, it is fitted to data obtained from mice subjected to a day/night cycle, providing a natural parameterization of its limit cycle. The origin of circadian time for the model was thus chosen to coincide with Zeitgeber time 0 in experiments (27, 28), allowing us to examine the obtained PRC from the perspective of synchronization with the diurnal cycle.

Fig. 5 displays a catalog of the PRC associated to most parameters of the AIR model, these PRC being obtained by applying a 10% increase in the corresponding parameter for one hour (blue curve) or 12 hours (red curve). The response to the one-hour stimulus provides us with information about the instantaeous sensitivity of the clock to the corresponding input throughout the circadian cycle. The 12-hour stimulus protocol is relevant for understanding the alternation between contrasting metabolic phases lasting 12 hours. For example, the value of the 12-hour PRC at time 0 characterizes the exposure to the stimulus between 0 and 12 hours (i.e., the subjective day). The intervals of subjective day (inactive period) and night (active period) in the experiments are indicated as white and gray regions in Fig. 5.

**Figure 5.**
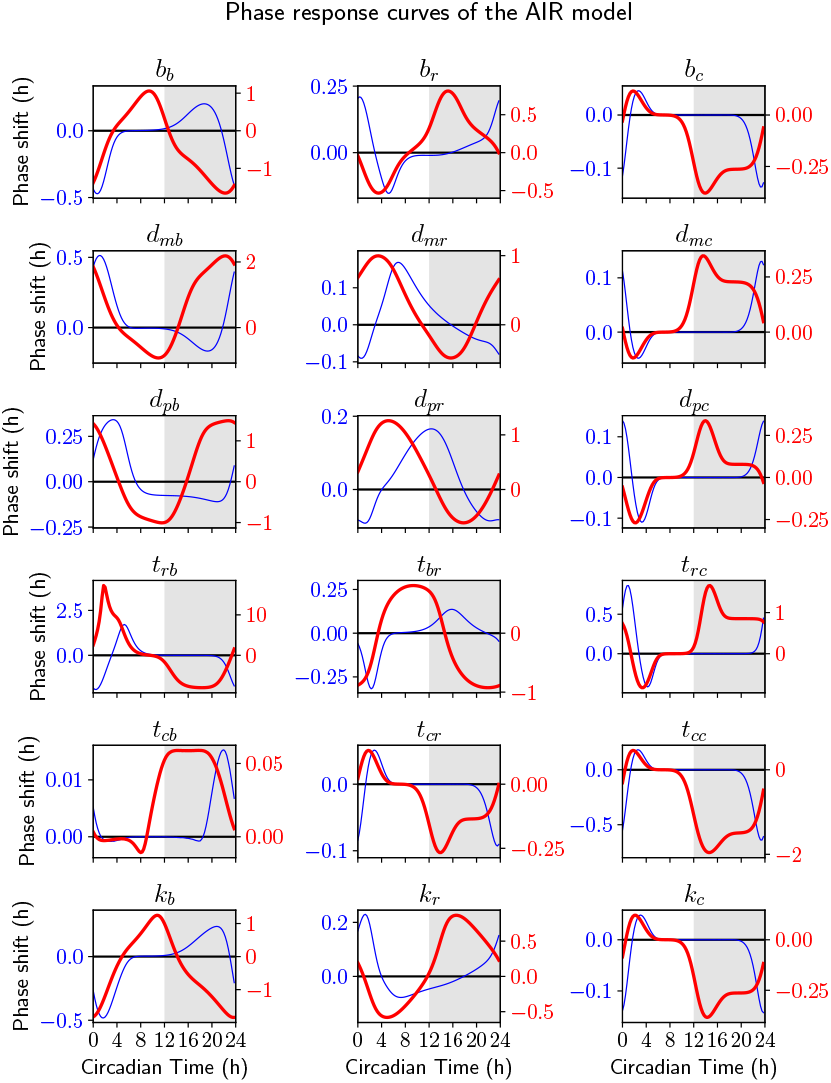
Phase response curves of the AIR model. From left to right, each of the three columns corresponds to mechanisms where respectively *Bmal1, Reverb*, or *Cry* is the main actor. Blue curves are obtained with a 1-hour stimulus starting at the given time, while the red ones characterize 12-hour stimuli.

Remarkably, Fig. 5 indicates that the modulation of *t*_*rb*_ (i.e., the threshold of regulation of *Reverb* by BMAL, defined as the BMAL concentration at which *Reverb* is half-activated, for zero CRY concentration), stands out as the most potent input, both for the 1-hour and the 12-hour protocol. This correlates with this input being a strong modifier of the limit cycle.

Besides the major input discussed above, the largest effects were here also observed in response to the modulation of the parameters *b*_*b*_, *d*_*mb*_, *d* _*pb*_ and *k*_*b*_ (Fig. 5, first column), which directly govern BMAL abundance and were also identified in connection with limit cycle changes (Table 3). Other interesting mechanisms are those destabilizing *Reverb* mRNA or REVERB protein (kinetic constants *d*_*mr*_ and *d* _*pr*_).

In comparison, the CRY-related inputs gave rise to relatively modest phase shifts, with perhaps the exception of the *t*_*cc*_ input, which would correspond to the modulation of *Cry* transcription by its own protein. This is a surprising result since there are many reports on inputs targeting the biological CRYs or PERs (both represented by CRY in the model) (see, e.g., (37, 38)), as we shall discuss below.

Because a change in oscillation period naturally induces a phase shift and may thus contribute to phase resetting mechanisms, we also computed how this period depends on kinetic constants (Fig. S1). Since we considered a wider range of variations, Fig. S1 provides information on potential phase resetting which is complementary to Fig. 5.

### Plausible synchronization mechanisms

To be relevant for synchronization, input mechanisms must not only correct a period mismatch between the clock and the day/night cycle (39), but also maintain a stable phase relationship between them. This puts constraints on the mechanisms informing the clock of the time of the day.

For simplicity of discussion, we will assume here that metabolic stimuli are aligned with the day/night alternation and are associated with a single clock parameter, with this parameter taking one value during the day and another during the night.

The liver clock in vivo has a natural period close to 24 hours (40), and thus does not require a daily phase shift. A physiological stimulus should thus begin when the PRC crosses zero (35), and under the above assumption, this zero should be located near dawn or dusk. To ensure stability, a negative phase correction should be applied when the clock is ahead, since the stimulus will then start at a larger phase than normal. Since the PRCs in Fig. 5 were computed for a positive modulation the kinetic constant, the slope of the PRC at the zero should be negative (resp., positive) if the studied stimulus increases (resp., decreases) this constant.

We thus examined Fig. 5 for PRCs statisfying the above criteria, since inputs producing such PRCs would, under our simplifying assumptions, ensure synchronization of the clock model to a day/night metabolic alternation with approximatively the correct peak timings. We identified the following mechanisms:

1. The affinity of BMAL1 for DNA is decreased (*t*_*rb*_ increased) during the night (*t*_*rb*_ panel, with PRC zero near CT 9.5). This is consistent with the NAD+-dependent protein deacetylase SIRT1 decreasing the transcription factor activity of the BMAL1:CLOCK complex (33, 41), given that NAD+ levels are on average higher during the night (data from (42) shown in (8)). Note that phosporylation of BMAL1 and CLOCK also influences the DNA binding of CLOCK:BMAL1 (43–45).
2. The *Bmal1* basal transcription rate is increased during the night (*b*_*b*_ panel, with PRC zero near ZT 12.75). A strong modulator of this rate is the PGC1*α* protein, which is indeed nuclear during the night (34). Numerical simulations by Woller et al. (8) showed that modulation of *Bmal1* expression by the AMPK-PGC1*α*-BMAL1 axis reinforced clock oscillations, which was confirmed by Foteinou et al. (9).
3. The REVERB protein is destabilized during the night (*d* _*pr*_ panel, with PRC zero near ZT 13.25). REVERB*α* is stabilized by GSK3*β*, whose inhibition by lithium leads to a rapid degradation of REVERB*α*(46). Besing et al. (47) found that in the SCN, GSK3*β* is mostly phosphorylated and active between ZT2 and ZT14 (i.e, during the day).
4. The CRY protein is destabilized during the day (*d* _*pc*_ panel, with PRC zero near ZT 23.25). Lamia et al. (37) showed that CRY degradation is induced by AMPK activity, which is indeed higher during the day in the liver (see profiles in (8), compatible with data in (42)).

An important synchronizing mechanism missing in this list is the upregulation of both PER transcription and translation by insulin and Insulin Growth Factor 1 (38). In our minimal model, this would correspond to a positive modulation of *b*_*c*_ and of *k*_*c*_, since CRY represents both PERs and CRYs. The PRCs for these two parameters are essentially identical (third column of Fig. 5, top and bottom row), and feature a negative-slope zero around CT8, which should thus be the beginning of a 12-hour stimulus to achieve stable entrainment. Now, Mukherji et al. (48) observed that insulin level is minimal at ZT4 and maximal at ZT16, and is thus above average between ZT10 and ZT22, in good agreement with the optimal window mentioned above. Thus, it appears that for the insulin pathway also, the predictions of our simple model are quite consistent with experimental observations.

## DISCUSSION

In this work, we designed a minimal mathematical model of the mammalian circadian clock that reproduces surprinsingly well (Fig. 2) the experimental expression profiles obtained by Hughes et al. (27) and Narumi et al. (28) from mouse livers. This model is based on three actors *Bmal1, Reverb*, and *Cry*, each of them playing a different mechanistic role: *Bmal1* activates the two other genes, *Reverb* represses the two other genes, and *Cry* inhibits the induction by *Bmal1* of the two other genes. Therefore, we named this model the Activator-Inhibitor-Repressor model (AIR) model. Such simple models can prove useful to better understand the design principles of the circadian clock and form a base on which to build more refined models. Note that a similar structure was described in a reduced model obtained by Almeida et al. (49), although without the repression of *Cry* by REVERB.

This model was obtained through a systematic reduction of a more complex model. To assess the relevance of an actor, we assessed whether its temporal variation, rather than its mere presence, was necessary to reproduce the observed clock dynamics. This is in line with recent experiments which tested the role of a molecular actor not by knocking it out, but by making its action constant in time. For example, Matsumura et al. (50) designed mutant mouse lines where expression of Cry1 and Cry2 was maintained but was made constant (50). Similarly, Abe et al. abolished the transcriptional rhythms of *Bmal1* by deleting the RRE in its promoter (44). If the influence of an actor can be made constant without perturbing the clock gene expression profiles, we can conclude that it is not essential to the clock dynamics. Theoretically, this approach was pionereed by Pett et al. (24).

Numerically, the situation is different, since control parameters are generally not known but are optimized to fit the model to the data. However, similar studies can be carried out by progressively flattening the time profile of a molecular actor while simulteously maintaining the agreement of the mathematical model with the data.

Moreover, we found that a two-gene model featuring only *Bmal1* and *Reverb* captured the experimental data quite well. This suggests that these two genes have their own dynamics and that the negative feedback loop formed by these two genes is the primary generator of clock oscillations. This is consistent with reports that knocking down *Reverbα, β* (51–53) severely disrupts the clock. Closer to the methodology followed here, Matsumura et al. suppressed oscillations in *Cry1* and *Cry2* expression, and found that oscillations in *Bmal1* and *RevErbα* expression were almost unchanged, being merely phase-delayed (50). Moreover, Fougeray et al. observed that knocking out the insulin receptor barely modified the *Bmal1* and *Reverbα* profiles (54). Interestingly, such a simple activator-repressor loop is at the heart of the circadian clock of *Ostreococcus tauri*, the smallest eukaryote known (55, 56).

Trying to suppress BMAL1 oscillations in the three-gene model significantly degraded fit quality. This is to be contrasted with the recent observation that abolishing the transcriptional rhythms of *Bmal1* by deleting the RRE in its promoter does not suppress circadian oscillations, making them only less robust to perturbations (44). In this study, however, oscillations in the phosphorylation of BMAL1 persisted as well as oscillations in *Clock* expression, suggesting that the activity of the CLOCK:BMAL1 complex was still varying in time. Therefore, these results are not contradictory with our finding that variations of BMAL1 expression play an important role, since in our model, the protein BMAL1 actually describes the activity of the CLOCK:BMAL1 complex.

We also found that taking into account the repression of *Cry* by REVERB (31) is important to achieve quantitative agreement with experimental data, as previously noted (7, 8). Importantly, this interaction induces a positive feedback loop where CRY inhibits *Reverb* transcription and REVERB represses *Cry* transcription.

Moreover, we totally lost oscillations when we tried to suppress the cycling of REVERB abundance, suggesting that REVERB is the central actor for generating the circadian oscillations. Interestingly, REVERB is at the intersection of the BMAL-REVERB negative feedback loop and the CRY-REVERB positive feedback loop. Such a combination of positive and negative feedback loops has been shown to be essential for generating robust oscillations in biological oscillators (57, 58).

In a similar study, Pett et al. (24) obtained another three-gene loop model, featuring *Reverb, Per* and *Cry* arranged in a “repressilator” motif, where each gene represses another and is repressed by the third one. However, their model assumed that PER and CRY are repressors of all genes except *Bmal1*, rather than inhibitors of BMAL1 inducing the same genes. In this hypothesis, the PER and CRY proteins can act independently of each other, and even in the absence of BMAL1. This may explain that they arrived at a different conclusion than the present study, since our model assumes that inhibition by PER and CRY depends on their simultaneous colocalization with the BMAL1:CLOCK complex (59). Moreover, Pett et al. (60), elaborating on (24), found that in liver, the BMAL1-REVERB loop is actually dominant compared to the repressilator loop, which is consistent with our findings.

Having obtained an AIR model reproducing precisely oscillations in mRNA and protein abundances of the liver clock, we characterized its sensitivity to perturbations, which provides complementary information to profile fitting. We first studied how the location of the limit cycle in the six-dimensional state space varied following a permanent perturbation of the control parameters. We found that the limit cycle was most sensitive to variations in the transcriptional activity of BMAL1, consistent with its known modulation by SIRT1 (33), and in the abundance of BMAL1, consistent with the control of *Bmal1* transcription by PGC1*α* (8, 9).

We then computed the phase response curves (PRC) characterizing the phase shift following a transient modification of a control parameter depending on the time of the perturbation, for 1-hour and 12-hour stimuli. The former provide us with information about the instantaneous response of the clock at different moments of the cycle and the latter about the reaction to metabolic phases during 12 hours. Since PRCs are very sensitive to changes in the molecular circuitry, comparing experimental and theoretical PRCs is a stringent test for mathematical models (61, 62) and may provide hints about the input mechanisms. Note that the two types of sensitivity are not unrelated, since a large change in the limit cycle is prone to induce a large phase shift. Given experimentally measured PRC, a future extension of this work could be to obtain a reduced model preserving the PRC, as done by Taylor et al. (23).

Remarkably, we found that the model-predicted behavior of a few mechanisms could be related to established mechanisms driving the circadian clock. These were in particular: (1) the modulation of the transcription factor activity of BMAL1, such as induced by SIRT1 (33, 41); (2) the modulation of *Bmal1* transcription, such as induced by PGC1*α* (8, 9, 34); (3) the modulation of CRY1 stability, such as controlled by AMPK activity (37); (4) the modulation of REVERB*α* stability such as governed by GSK3*β* (47, 63); (5) the insulin-induced upregulation of PER mRNA and protein (38).

The qualitative agreement between model predictions and mechanisms from the literature is quite surprising given the simplicity of the model and the fact that real input pathways are often gated (64), being active only in temporal windows whose position and duration are optimized with respect to clock function (see, e.g., (55, 65)). Moreover, resetting is generally achieved through multiple signals, and each of these signals acts through several pathways. For example, AMPK stimulates CRY1 degradation (37) but also activates PGC1*α* (66) and thus *Bmal1* expression. Insulin upregulates PER (38) but also reduces PGC1*α* expression levels (67). The present work is thus only the first step towards more detailed studies.

## CONCLUSION

We designed a very simple model fitting well data for the hepatic clock, and whose clear design may help to understand better the role of the different clock genes but also to analyze experimental results. In particular we found that *Bmal1* and *Reverb* seem important to generate the oscillation profiles while *Cry* may play an important role for clock resetting. The phase response curves predicted by this model are compatible with a number of already identified entrainment mechanisms, which is quite satisfying given the simplicity of the model.

## AUTHOR CONTRIBUTIONS

P.D. and M.L. designed and carried out the research, and wrote the paper.

## ACKNOWLEDGMENTS

The authors acknowledge funding support from ANR through LabEx CEMPI (ANR-11-LABX-0007).

## Supplemental information

The three-gene AIR model (BMAL1/REVERB/CRY) has been deposited in the BioModels database under accession MODEL2511090001 and will be available at https://www.ebi.ac.uk/biomodels/MODEL2511090001 after final publication.

The two-gene AR model (BMAL1/REVERB) has been deposited in the BioModels database under accession MODEL2511090002 and will be available at https://www.ebi.ac.uk/biomodels/MODEL2511090002 after final publication.

**Figure S1.**
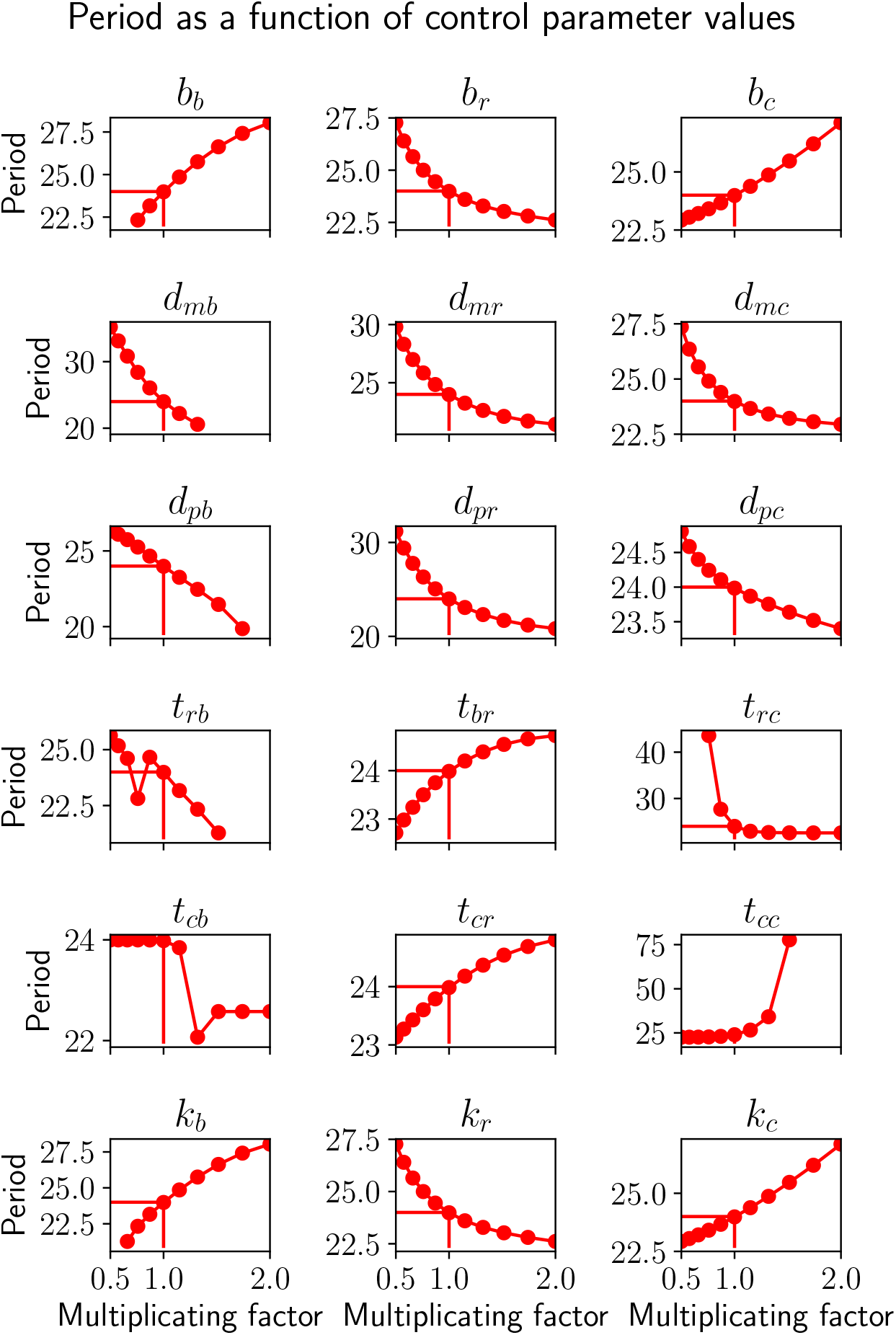
Variation of the period of the AIR Model with control parameter values. Only the parameter values for which the clock amplitude was above half of the nominal value are shown. The period is always of 24 hours for the nominal parameter value.

**Table S1.**
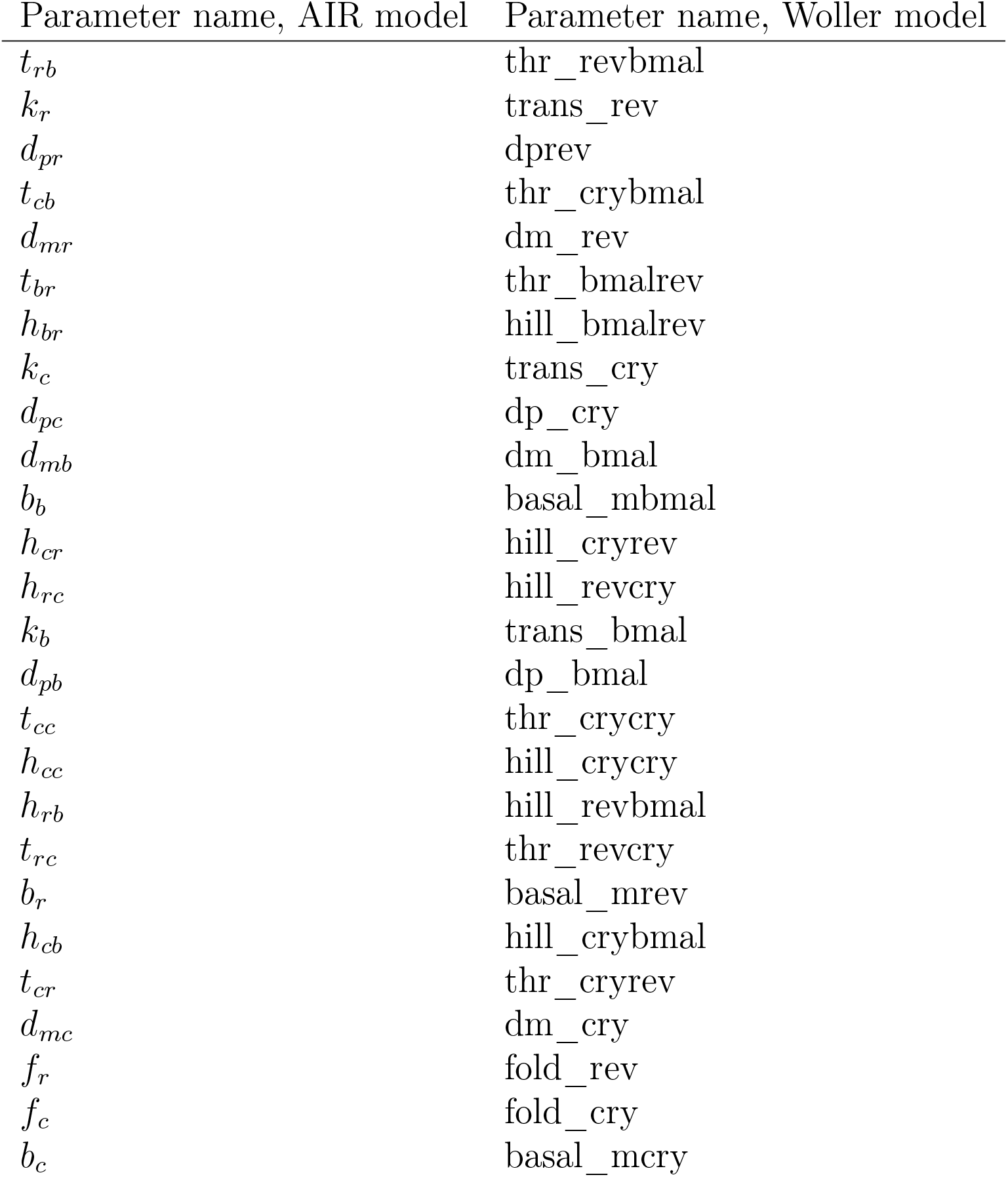
Correspondence between parameter names in the AIR model and in the Woller model.

## REFERENCES

1. Dibner, C., U. Schibler, and U. Albrecht, 2010. The mammalian circadian timing system: organization and coordination of central and peripheral clocks. Annu Rev Physiol 72:517–549. 10.1146/annurev-physiol-021909-135821.

2. Takahashi, J. S., 2017. Transcriptional architecture of the mammalian circadian clock. Nat. Rev. Genet. 18:164– 179.

3. Forger, D. B., and C. S. Peskin, 2003. A detailed predictive model of the mammalian circadian clock. Proc Natl Acad Sci U S A 100:14806–14811. 10.1073/pnas.2036281100.

4. Leloup, J.-C., and A. Goldbeter, 2003. Toward a detailed computational model for the mammalian circadian clock. Proc Natl Acad Sci U S A 100:7051–7056. 10.1073/pnas.1132112100.

5. Becker-Weimann, S., J. Wolf, H. Herzel, and A. Kramer, 2004. Modeling feedback loops of the Mammalian circadian oscillator. Biophys J 87:3023–3034. 10.1529/biophysj.104.040824.

6. Mirsky, H. P., A. C. Liu, D. K. Welsh, S. A. Kay, and F. J. Doyle, 2009. A model of the cell-autonomous mammalian circadian clock. Proc Natl Acad Sci U S A 106:11107–11112. 10.1073/pnas.0904837106.

7. Relógio, A., P. O. Westermark, T. Wallach, K. Schellenberg, A. Kramer, and H. Herzel, 2011. Tuning the mammalian circadian clock: robust synergy of two loops. PLoS Comput Biol 7:e1002309. 10.1371/journal.pcbi.1002309.

8. Woller, A., H. Duez, B. Staels, and M. Lefranc, 2016. A Mathematical Model of the Liver Circadian Clock Linking Feeding and Fasting Cycles to Clock Function. Cell Rep 17:1087–1097. 10.1016/j.celrep.2016.09.060.

9. Foteinou, P. T., A. Venkataraman, L. J. Francey, R. C. Anafi, J. B. Hogenesch, and F. J. Doyle, 2018. Computational and experimental insights into the circadian effects of SIRT1. Proc Natl Acad Sci U S A 115:11643–11648.

10. Leloup, J.-C., and A. Goldbeter, 2004. Modeling the mammalian circadian clock: sensitivity analysis and multiplicity of oscillatory mechanisms. J. Theor. Biol. 230:541– 562.

11. Stelling, J., E. D. Gilles, and F. J. Doyle, 2004. Robustness properties of circadian clock architectures. Proc. Natl. Acad. Sci. U. S. A. 101:13210–13215.

12. Zeilinger, M. N., E.M. Farré, S. R. Taylor, S. A. Kay, and F. J. Doyle, 2006. A novel computational model of the circadian clock in Arabidopsis that incorporates PRR7 and PRR9. Mol. Syst. Biol. 2:58.

13. Pfeuty, B., J.-F. Bodart, R. Blossey, and M. Lefranc, 2012. A dynamical model of oocyte maturation unveils precisely orchestrated meiotic decisions. PLoS Comput. Biol. 8:e1002329.

14. Raue, A., C. Kreutz, T. Maiwald, J. Bachmann, M. Schilling, U. Klingmüller, and J. Timmer, 2009. Structural and practical identifiability analysis of partially observed dynamical models by exploiting the profile likelihood. Bioinformatics 25:1923–1929.

15. Raue, A., V. Becker, U. Klingmüller, and J. Timmer, 2010. Identifiability and observability analysis for experimental design in nonlinear dynamical models. Chaos 20:045105.

16. St John, P. C., and F. J. Doyle, 2013. Estimating confidence intervals in predicted responses for oscillatory biological models. BMC Syst. Biol. 7:71.

17. Jolley, C. C., M. Ukai-Tadenuma, D. Perrin, and H. R. Ueda, 2014. A mammalian circadian clock model incorporating daytime expression elements. Biophys J 107:1462–1473. 10.1016/j.bpj.2014.07.022.

18. Apgar, J. F., D. K. Witmer, F. M. White, and B. Tidor, 2010. Sloppy models, parameter uncertainty, and the role of experimental design. Mol. Biosyst. 6:1890–1900.

19. Gutenkunst, R. N., J. J. Waterfall, F. P. Casey, K. S. Brown, C. R. Myers, and J. P. Sethna, 2007. Universally sloppy parameter sensitivities in systems biology models. PLoS Comput. Biol. 3:1871–1878.

20. Kim, J. K., and D. B. Forger, 2012. A mechanism for robust circadian timekeeping via stoichiometric balance. Mol. Syst. Biol. 8:630.

21. Clodong, S., U. Dühring, L. Kronk, A. Wilde, I. Axmann, H. Herzel, and M. Kollmann, 2007. Functioning and robustness of a bacterial circadian clock. Molecular systems biology 3:90.

22. Del Olmo, M., A. Kramer, and H. Herzel, 2019. A Robust Model for Circadian Redox Oscillations. Int. J. Mol. Sci. 20.

23. Taylor, S. R., F. J. Doyle, and L. R. Petzold, 2008. Oscillator model reduction preserving the phase response: application to the circadian clock. Biophy. J 95:1658– 1673.

24. Pett, J. P., A. Korenčič, F. Wesener, A. Kramer, and H. Herzel, 2016. Feedback Loops of the Mammalian Circadian Clock Constitute Repressilator. PLoS Comput. Biol. 12:e1005266.

25. Hoops, S., S. Sahle, R. Gauges, C. Lee, J. Pahle, N. Simus, M. Singhal, L. Xu, P. Mendes, and U. Kummer, 2006. COPASI–a COmplex PAthway SImulator. Bioinformatics 22:3067–3074.

26. Bergmann, F., 2023. BASICO: A simplified Python interface to COPASI. JOSS 8:5553.

27. Hughes, M. E., L. DiTacchio, K. R. Hayes, C. Vollmers, S. Pulivarthy, J. E. Baggs, S. Panda, and J. B. Hogenesch, 2009. Harmonics of circadian gene transcription in mammals. PLoS Genet 5:e1000442. 10.1371/journal.pgen.1000442.

28. Narumi, R., Y. Shimizu, M. Ukai-Tadenuma, K. L. Ode, G. N. Kanda, Y. Shinohara, A. Sato, K. Matsumoto, and H. R. Ueda, 2016. Mass spectrometry-based absolute quantification reveals rhythmic variation of mouse circadian clock proteins. Proc Natl Acad Sci U S A 113:E3461–E3467. 10.1073/pnas.1603799113.

29. Menet, J. S., S. Pescatore, and M. Rosbash, 2014. CLOCK:BMAL1 is a pioneer-like transcription factor. Genes Dev. 28:8–13. 10.1101/gad.228536.113.

30. Gonze, D., and W. Abou-Jaoudé, 2013. The Goodwin model: behind the Hill function. PLoS One 8:e69573.

31. Liu, A. C., H. G. Tran, E. E. Zhang, A. A. Priest, D. K. Welsh, and S. A. Kay, 2008. Redundant function of REV-ERBalpha and beta and non-essential role for Bmal1 cycling in transcriptional regulation of intracellular circadian rhythms. PLoS Genet 4:e1000023. 10.1371/journal.pgen.1000023.

32. Yin, L., and M. A. Lazar, 2005. The orphan nuclear receptor Rev-erbalpha recruits the N-CoR/histone deacetylase 3 corepressor to regulate the circadian Bmal1 gene. Mol. Endocrinol. 19:1452–1459.

33. Nakahata, Y., M. Kaluzova, B. Grimaldi, S. Sahar, J. Hirayama, D. Chen, L. P. Guarente, and P. Sassone-Corsi, 2008. The NAD+-dependent deacetylase SIRT1 modulates CLOCK-mediated chromatin remodeling and circadian control. Cell 134:329–340. 10.1016/j.cell.2008.07.002.

34. Liu, C., S. Li, T. Liu, J. Borjigin, and J. D. Lin, 2007. Transcriptional coactivator PGC-1alpha integrates the mammalian clock and energy metabolism. Nature 447:477–481. 10.1038/nature05767.

35. Forger, D. B., 2017. Biological Clocks, Rhythms, and Oscillations: The Theory of Biological Timekeeping. MIT Press.

36. Pfeuty, B., Q. Thommen, and M. Lefranc, 2011. Robust entrainment of circadian oscillators requires specific phase response curves. Biophys. J 100:2557–2565.

37. Lamia, K. A., U. M. Sachdeva, L. DiTacchio, E. C. Williams, J. G. Alvarez, D. F. Egan, D. S. Vasquez, H. Juguilon, S. Panda, R. J. Shaw, C. B. Thompson, and R. M. Evans, 2009. AMPK regulates the circadian clock by cryptochrome phosphorylation and degradation. Science 326:437–440. 10.1126/science.1172156.

38. Crosby, P., R. Hamnett, M. Putker, N. P. Hoyle, M. Reed, C. J. Karam, E. S. Maywood, A. Stangherlin, J. E. Chesham, E. A. Hayter, L. Rosenbrier-Ribeiro, P. Newham, H. Clevers, D. A. Bechtold, and J. S. O’Neill, 2019. Insulin/IGF-1 Drives PERIOD Synthesis to Entrain Circadian Rhythms with Feeding Time. Cell 177:896–909.e20.

39. Bordyugov, G., U. Abraham, A. Granada, P. Rose, K. Imkeller, A. Kramer, and H. Herzel, 2015. Tuning the phase of circadian entrainment. J. R. Soc. Interface 12:20150282.

40. Sinturel, F., P. Gos, V. Petrenko, C. Hagedorn, F. Kreppel, K.-F. Storch, D. Knutti, A. Liani, C. Weitz, Y. Emmenegger, P. Franken, L. Bonacina, C. Dibner, and U. Schibler, 2021. Circadian hepatocyte clocks keep synchrony in the absence of a master pacemaker in the suprachiasmatic nucleus or other extrahepatic clocks. Genes Dev. 35:329–334.

41. Bellet, M. M., Y. Nakahata, M. Boudjelal, E. Watts, D. E. Mossakowska, K. A. Edwards, M. Cervantes, G. Astarita, C. Loh, J. L. Ellis, G. P. Vlasuk, and P. Sassone-Corsi, 2013. Pharmacological modulation of circadian rhythms by synthetic activators of the deacetylase SIRT1. Proc Natl Acad Sci U S A 110:3333–3338. 10.1073/pnas.1214266110.

42. Hatori, M., C. Vollmers, A. Zarrinpar, L. DiTacchio, E. A. Bushong, S. Gill, M. Leblanc, A. Chaix, M. Joens, J. A. J. Fitzpatrick, M. H. Ellisman, and S. Panda, 2012. Time-restricted feeding without reducing caloric intake prevents metabolic diseases in mice fed a high-fat diet. Cell Metab 15:848–860. 10.1016/j.cmet.2012.04.019.

43. Brenna, A., and U. Albrecht, 2020. Phosphorylation and Circadian Molecular Timing. Front. Physiol. 11:612510.

44. Abe, Y. O., H. Yoshitane, D. W. Kim, S. Kawakami, M. Koebis, K. Nakao, A. Aiba, J. K. Kim, and Y. Fukada, 2022. Rhythmic transcription of Bmal1 stabilizes the circadian timekeeping system in mammals. Nat. Commun. 13:4652.

45. Otobe, Y., E. M. Jeong, S. Ito, Y. Shinohara, N. Kurabayashi, A. Aiba, Y. Fukada, J. K. Kim, and H. Yoshitane, 2024. Phosphorylation of DNA-binding domains of CLOCK-BMAL1 complex for PER-dependent inhibition in circadian clock of mammalian cells. Proc. Natl. Acad. Sci. U. S. A. 121:e2316858121.

46. Yin, L., J. Wang, P. S. Klein, and M. A. Lazar, 2006. Nuclear receptor Rev-erbalpha is a critical lithium-sensitive component of the circadian clock. Science 311:1002– 1005.

47. Besing, R. C., J. R. Paul, L. M. Hablitz, C. O. Rogers, R. L. Johnson, M. E. Young, and K. L. Gamble, 2015. Circadian rhythmicity of active GSK3 isoforms modulates molecular clock gene rhythms in the suprachiasmatic nucleus. J Biol. Rhythms 30:155–160.

48. Mukherji, A., A. Kobiita, and P. Chambon, 2015. Shifting the feeding of mice to the rest phase creates metabolic alterations, which, on their own, shift the peripheral circadian clocks by 12 hours. Proc Natl Acad Sci U S A 112:E6683–E6690. 10.1073/pnas.1519735112.

49. Almeida, S., M. Chaves, and F. Delaunay, 2020. Control of synchronization ratios in clock/cell cycle coupling by growth factors and glucocorticoids. R. Soc. Open Sci. 7:192054.

50. Matsumura, R., K. Yoshimi, Y. Sawai, N. Yasumune, K. Kajihara, T. Maejima, T. Koide, K. Node, and M. Akashi, 2022. The role of cell-autonomous circadian oscillation of Cry transcription in circadian rhythm generation. Cell Rep. 39:110703.

51. Bugge, A., D. Feng, L. J. Everett, E. R. Briggs, S. E. Mullican, F. Wang, J. Jager, and M. A. Lazar, 2012. Rev-erbα and Rev-erbβ coordinately protect the circadian clock and normal metabolic function. Genes Dev 26:657–667. 10.1101/gad.186858.112.

52. Cho, H., X. Zhao, M. Hatori, R. T. Yu, G. D. Barish, M. T. Lam, L.-W. Chong, L. DiTacchio, A. R. Atkins, C. K. Glass, C. Liddle, J. Auwerx, M. Downes, S. Panda, and R. M. Evans, 2012. Regulation of circadian behaviour and metabolism by REV-ERB-alpha and REV-ERB-beta. Nature 485:123–127. 10.1038/nature11048.

53. Solt, L. A., Y. Wang, S. Banerjee, T. Hughes, D. J. Kojetin, T. Lundasen, Y. Shin, J. Liu, M. D. Cameron, R. Noel, S.-H. Yoo, J. S. Takahashi, A. A. Butler, T. M. Kamenecka, and T. P. Burris, 2012. Regulation of circadian behaviour and metabolism by synthetic REV-ERB agonists. Nature 485:62–68. 10.1038/nature11030.

54. Fougeray, T., A. Polizzi, M. Régnier, A. Fougerat, S. Ellero-Simatos, Y. Lippi, S. Smati, F. Lasserre, B. Tramunt, M. Huillet, L. Dopavogui, J. Salvi, E. Nédélec, V. Gigot, L. Smith, C. Naylies, C. Sommer, J. T. Haas, W. Wahli, H. Duez, P. Gourdy, L. Gamet-Payrastre, A. Benani, A.-F. Burnol, N. Loiseau, C. Postic, A. Montagner, and H. Guillou, 2022. The hepatocyte insulin receptor is required to program the liver clock and rhythmic gene expression. Cell Rep. 39:110674.

55. Thommen, Q., B. Pfeuty, P.-E. Morant, F. Corellou, F.-Y. Bouget, and M. Lefranc, 2010. Robustness of circadian clocks to daylight fluctuations: hints from the picoeucaryote Ostreococcus tauri. PLoS Comput. Biol. 6:e1000990.

56. Morant, P.-E., Q. Thommen, B. Pfeuty, C. Vandermoere, F. Corellou, F.-Y. Bouget, and M. Lefranc, 2010. A robust two-gene oscillator at the core of Ostreococcus tauri circadian clock. Chaos 20:045108.

57. Ferrell, J. E., T. Y.-C. Tsai, and Q. Yang, 2011. Modeling the cell cycle: why do certain circuits oscillate? Cell 144:874–885.

58. Tsai, T. Y.-C., Y. S. Choi, W. Ma, J. R. Pomerening, C. Tang, and J. E. Ferrell, 2008. Robust, tunable biological oscillations from interlinked positive and negative feedback loops. Science 321:126–129.

59. Ye, R., C. P. Selby, Y.-Y. Chiou, I. Ozkan-Dagliyan, S. Gaddameedhi, and A. Sancar, 2014. Dual modes of CLOCK:BMAL1 inhibition mediated by Cryptochrome and Period proteins in the mammalian circadian clock. Genes Dev. 28:1989–1998.

60. Pett, J. P., M. Kondoff, G. Bordyugov, A. Kramer, and H. Herzel, 2018. Co-existing feedback loops generate tissue-specific circadian rhythms. Life Sci. Alliance 1:e201800078.

61. Taylor, S. R., A. Cheever, and S. M. Harmon, 2014. Velocity response curves demonstrate the complexity of modeling entrainable clocks. J Theor. Biol. 363:307–317.

62. Thommen, Q., B. Pfeuty, P. Schatt, A. Bijoux, F.-Y. Bouget, and M. Lefranc, 2015. Probing entrainment of Ostreococcus tauri circadian clock by green and blue light through a mathematical modeling approach. Front. Genet. 6:65.

63. Yin, L., N. Wu, J. C. Curtin, M. Qatanani, N. R. Szwergold, R. A. Reid, G. M. Waitt, D. J. Parks, K. H. Pearce, G. B. Wisely, and M. A. Lazar, 2007. Rev-erbalpha, a heme sensor that coordinates metabolic and circadian pathways. Science 318:1786–1789.

64. Geier, F., S. Becker-Weimann, A. Kramer, and H. Herzel, 2005. Entrainment in a model of the mammalian circadian oscillator. J. Biol. Rhythms 20:83–93.

65. Pfeuty, B., Q. Thommen, F. Corellou, E. B. Djouani-Tahri, F.-Y. Bouget, and M. Lefranc, 2012. Circadian clocks in changing weather and seasons: lessons from the picoalga Ostreococcus tauri. Bioessays 34:781–790.

66. Cantó, C., L. Q. Jiang, A. S. Deshmukh, C. Mataki, A. Coste, M. Lagouge, J. R. Zierath, and J. Auwerx, 2010. Interdependence of AMPK and SIRT1 for metabolic adaptation to fasting and exercise in skeletal muscle. Cell Metab 11:213–219. 10.1016/j.cmet.2010.02.006.

67. Herzig, S., F. Long, U. S. Jhala, S. Hedrick, R. Quinn, A. Bauer, D. Rudolph, G. Schutz, C. Yoon, P. Puigserver, A. Spiegelman, and M. Montminy, 2001. CREB regulates hepatic gluconeogenesis through the coactivator PGC-1. Nature 413:179–183.

