## Supplemental information for "A minimal activator-inhibitor-repressor model of the hepatic circadian clock"

November 9, 2025

The three-gene AIR model (BMAL1/REVERB/CRY) has been deposited in the BioModels database under accession MODEL2511090001 and will be available at `https://www.ebi.ac.uk/biomodels/MODEL2511090001` after final publication.

The two-gene AR model (BMAL1/REVERB) has been deposited in the BioModels database under accession MODEL2511090002 and will be available at `https://www.ebi.ac.uk/biomodels/MODEL2511090002` after final publication.

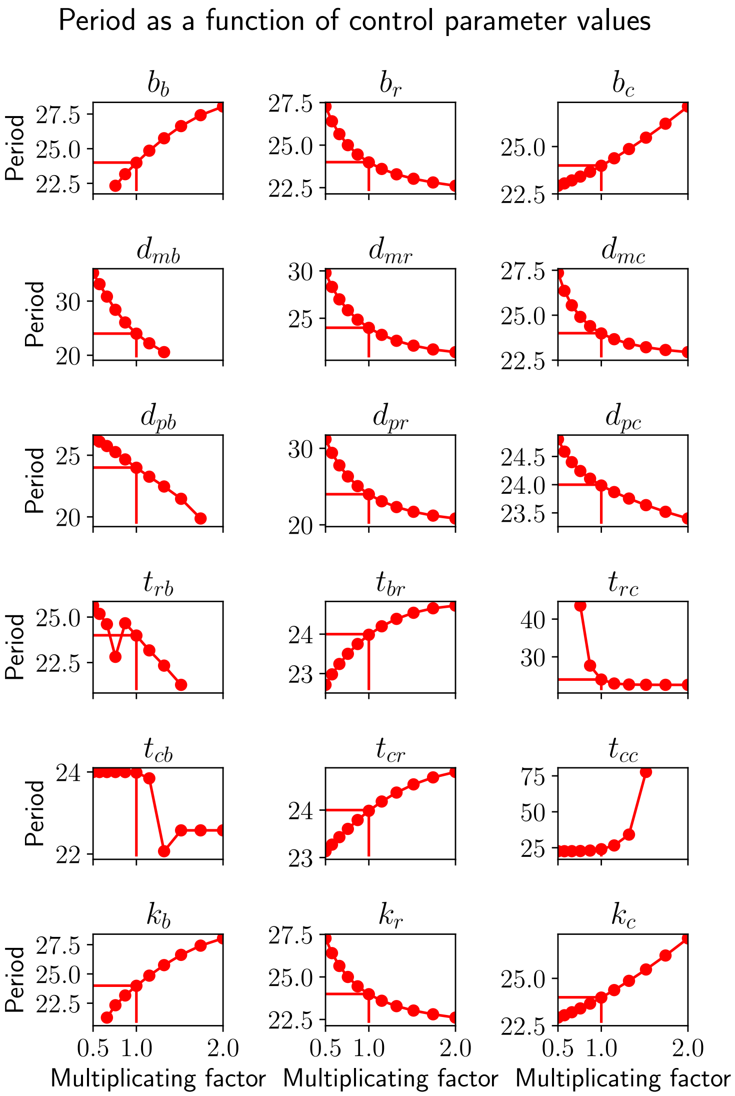


Figure S1: Variation of the period of the AIR Model with control parameter values. Only the parameter values for which the clock amplitude was above half of the nominal value are shown. The period is always of 24 hours for the nominal parameter value.

| Parameter name, AIR model | Parameter name, Woller model |
|---|---|
| $t_{rb}$ | thr_revbmal |
| $k_r$ | trans_rev |
| $d_{pr}$ | dprev |
| $t_{cb}$ | thr_crybmal |
| $d_{mr}$ | dm_rev |
| $t_{br}$ | thr_bmalrev |
| $h_{br}$ | hill_bmalrev |
| $k_c$ | trans_cry |
| $d_{pc}$ | dp_cry |
| $d_{mb}$ | dm_bmal |
| $b_b$ | basal_mbmal |
| $h_{cr}$ | hill_cryrev |
| $h_{rc}$ | hill_revcry |
| $k_b$ | trans_bmal |
| $d_{pb}$ | dp_bmal |
| $t_{cc}$ | thr_crycry |
| $h_{cc}$ | hill_crycry |
| $h_{rb}$ | hill_revbmal |
| $t_{rc}$ | thr_revcry |
| $b_r$ | basal_mrev |
| $h_{cb}$ | hill_crybmal |
| $t_{cr}$ | thr_cryrev |
| $d_{mc}$ | dm_cry |
| $f_r$ | fold_rev |
| $f_c$ | fold_cry |
| $b_c$ | basal_mcry |

Table S1: Correspondence between parameter names in the AIR model and in the Woller model
